# Microbial community of a colloidal activated carbon barrier at a perfluoroalkyl substance impacted site and sorption of perfluorooctanesulfonic acid by bacteria

**DOI:** 10.1101/2025.09.19.677121

**Authors:** Brian Eddie, Christopher Katilie, Lindsay Van House, Dagmar Leary, Lina Bird

## Abstract

Per- and polyfluorinated alkyl substances (PFAS) including perfluorooctanesulfonic acid (PFOS) are extremely refractory anthropogenic compounds that are widespread in the environment. Their fate and transport are of increasing interest, including the ways that biological activity may affect the efficiency of state-of-the-art mitigation efforts, such as colloidal activated carbon (CAC) injection. The microbial communities of sediments in and around a CAC barrier from 2.7 to 9 m below surface were investigated using 16S ribosomal RNA amplicon sequencing and strains isolated from the samples. The microbial community consisted of taxa commonly associated with the subsurface environment, including a high proportion of *Bathyarchaeota* in the deeper samples. The presence of PFAS or the CAC barrier used to treat it affected the microbial community. Bacteria in the order *Peptococcales* had higher relative abundance in the CAC barrier than upstream or downstream of it. The inverse Simpson index decreased with depth, indicating less variety in the microbes that were present, but was higher in the samples downstream of the CAC barrier, which may indicate an impact of the PFAS on the microbial community. The sorption of PFOS was investigated for six newly isolated strains as well as four other strains, and results indicate that sorption of PFOS by different species or genera of bacteria is more variable than previously thought. These results suggest that the microbial community could play a role in the mobility of PFAS within subsurface waters, and emphasizes the need to understand the subsurface microbial community and its interactions with PFAS.

## 1. Introduction

Per- and polyfluorinated alkyl substances (PFAS) are extremely refractory compounds that are widespread in the environment through a wide range of activities, including chemical production, agriculture, maintenance, and firefighting [1, 2]. Due to its use in aqueous film forming foams (AFFF) used in firefighting, perfluorooctanesulfonic acid (PFOS) and other PFAS are widespread contaminants in soils, aquifers, and water near firefighting training facilities. PFOS is regulated in many countries as potentially harmful at low concentrations, with drinking water limits for PFOS of 4 parts per trillion set to go into effect in the United States by 2029 [3]. Efforts are underway to prevent the spread of PFAS from point sources such as firefighting training sites, and understanding the factors that affect their migration through soil and sediment is relevant to efficiently designing systems to mitigate their spread [1].

The response of soil and sediment microbial communities to PFAS has been studied before [4], but these studies are typically limited to less than 2 m below the surface. Deeper samples from aquifers are typically limited to the groundwater and may not represent the biofilms present in these environments. PFAS have been reported to change the microbial communities, by increasing the proportion of *Proteobacteria* [5], decreasing the proportion of nitrifying bacteria [6] and suppressing *Dehalococcoides* with an increase in methanogenic archaea [7]. Additional studies help paint a more complete picture of PFAS impact on microbial communities, with some concentration dependent associations of the genera *Gordonia* and *Acidimicrobium* and negative correlations of the family *Oxalobacteraceae* to PFAS [8].

PFAS are accumulated to varying degrees by bacteria. Fitzgerald et al [9] showed that both the Gram-negative *Aliivibrio fischeri* and to a lesser extent the Gram-positive *Staphylococcus epidermidis* could partition various perfluorinated molecules, while Naumann et al showed the same with supported lipid monolayers [10], suggesting that the mechanism of PFOS partitioning in bacteria was its insertion into bacterial membranes. In contrast to the membrane insertion mechanism, Butzen et al [11] demonstrated pH dependent PFOS partitioning in *Bacillus subtilis* and *Pseudomonas putida*, with rapid sorption at pH 4 and desorption at pH 6, suggesting an electrostatic mechanism of adsorption. These studies, along with measurements of biofilms in the environment [12], suggested a generalizable rule that PFOS is sorbed by bacteria through multiple mechanisms.

We investigated the microbial community of a PFAS impacted site with an experimental colloidal activated carbon (CAC) barrier in the mid-Atlantic region of the US to identify changes in the microbial community inhabiting the subsurface upstream, within, and downstream of the barrier. This site was used for fire fighter training with AFFF, and had water concentrations of PFAS in the parts per billion range pre-treatment. Subsamples for DNA extraction and cultivation were taken during coring activities to monitor a CAC barrier one month after the barrier was established and two years later. As this *in situ* method of remediation is still in pilot phase, the effect of CAC on the microbial community has not previously been examined. We investigated the microbial community of the sediment to identify changes that may be associated with the CAC treatment area, and the presence of PFAS. Isolates were screened for ability to sorb PFOS. In our work, we demonstrate PFOS sorption varies even within closely related taxa. This indicates that the previous observations on PFOS sorption by bacteria are not universally generalizable.

## 2. Methods

### 2.1. Sample collection and bacterial isolation

Sediment samples were collected on September 13, 2022 and October 1 and 2, 2024 from direct push coring that was being conducted to evaluate the effectiveness of colloidal activated carbon (CAC) injections at a PFAS impacted site. These cores were taken upstream of or beside the barrier (US1, US4, US5, US10), in the barrier (IN2, IN3, IN6, IN7, IN8), and downstream of the barrier (DS9) (Figure 1A).

**Fig. 1.**
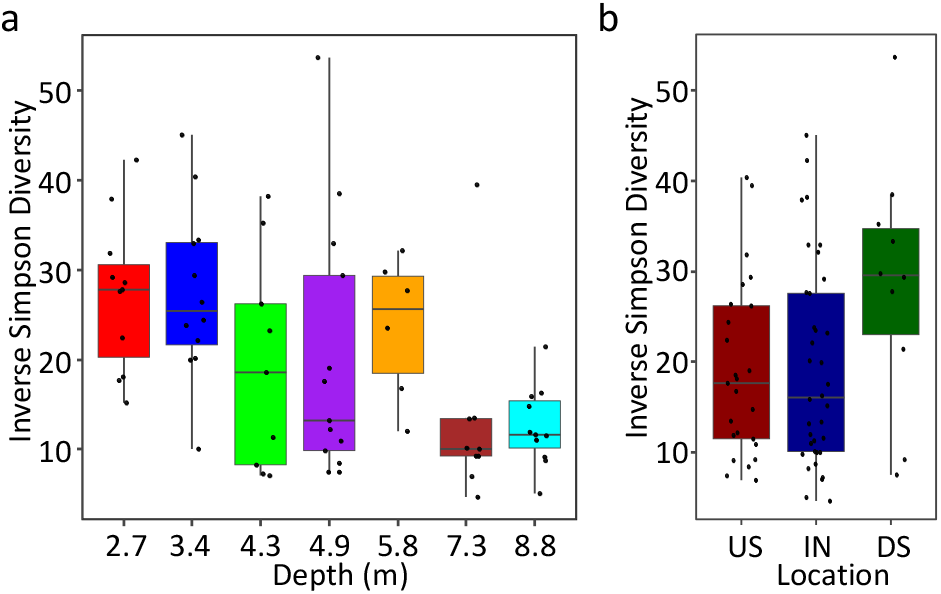
Inverse Simpson diversity metric. a) samples separated by depth below surface b) samples separated by location relative to the CAC barrier US – upstream, IN – in the barrier, DS – downstream of the barrier

Coring went to a depth of 9.1 m in 1.52 m sections. The cores were split lengthwise, and samples for DNA extraction were collected from 7 depths: 2.7 m, 3.4 m, 4.3 m, 4.9 m, 5.8 m, 7.3 m, and 8.8 m below the surface. A 50 mL centrifuge tube was used to scoop 20 - 50 g of sediment from the interior face of the split and stored on ice in 2022 and on dry ice in 2024 for transport back to the lab. Once there, they were stored at -80°C until extraction. Replicate samples were taken from US5 2.7 m, IN6 3.4 m, and DS9 4.9 m. Two field blank samples were collected in 2024 by pouring 5 mL of DNA- and nuclease-free water back and forth between two 50 mL centrifuge tubes three times.

Sediment and groundwater were collected in 2022 as starting material for enrichment cultures. A total of 500 µL of a 0.1 g/ml sediment/groundwater slurry was transferred to 5 mL of filter-sterilized groundwater spiked with 1 ppm perfluorooctanoic acid (PFOA) (Aldrich, St. Louis, MO, USA) and 1 g/L yeast extract. After shaking at 18°C for 1 week, cultures were transferred into Reasoner’s 2A liquid broth (R2A, Teknova, Hollister, CA, USA) spiked with 10 ppm PFOA. Following 1 week of shaking at 18°C, an aliquot of the culture was swabbed onto a 10 ppm PFOA and R2A agar plate and incubated at 30°C. After 24 h, the plate was speckled with opaque colonies that were pink or cream in color. Colonies were selected and swabbed onto fresh plates; this process was repeated until streaking produced uniform colonies for each strain. One colony for each strain was selected to inoculate fresh R2A, and the resulting culture was then aliquoted into 25% glycerol stocks frozen at −80°C. One strain, *Pseudomonas* sp. CBR-F, was isolated from granular activated carbon that had been used to filter PFAS impacted water using the same methods described above[13]. *Acinetobacter* sp. B1 was isolated by directly plating a 1:1000 dilution of sediment in R2A broth onto R2A agar supplemented with 10 ppm PFOA.

### 2.2. DNA extraction and library preparation

DNA extraction and library preparation were randomized into batches of 8 samples each to reduce systematic errors. Samples were thawed and mixed thoroughly by vortexing and shaking. DNA was extracted from a 0.5-0.8 g subsample using a Power Soil Pro kit (Qiagen, Germantown, MD, USA) following the manufacturer’s protocol with the following modifications. Lysis buffer was supplemented with 80 µL of ethylenediaminetetraacetic acid (EDTA) to aid in cell release from the sand grains. A second precipitation step was performed with an additional 100 µL of buffer CD2 to improve removal of amplification inhibitors. Two 100 µL aliquots of DNA/nuclease-free water and two 100 µL aliquots of the field blanks were extracted to control for DNA introduced by the kit, cross contamination, or poor barcode segregation. Samples from the first round of coring were processed and sequenced in June, 2024 and samples from the second round of coring were processed and sequenced in December 2024.

Amplicon libraries were generated by polymerase chain reaction (PCR) amplification of the variable region 4 (V4) of the 16S ribosomal RNA (rRNA) gene. Universal primers V4F-515Y-**ACACTCTTTCCCTACACGACGCTCTTCCGATCT**GTGYCAGCMGCCGCGGTAA and V4R-806RB – **GACTGGAGTTCAGACGTGTGCTCTTCCGATCT**GGACTACNVGGGTWTCTAAT that give good coverage of both bacteria and Archaea [14, 15] were used with an adapter sequence (bold) to allow for Amplicon-EZ sequencing (Azenta Biotech, Burlington, MA, USA). PCR reactions (50 µL total) used Q5 2X master mix (NEB, Ipswich, MA, USA), 400 nM each primer, and 1 µL of DNA extract as template. A spike in standard of 10 fg of amplified *Corynebacterium glutamicum* 16S rRNA gene (∼7000 copies) was added to the master mix to provide a baseline for comparison. In addition to the two field blanks and two extraction blank negative controls, one reaction with 1 ng of the spike-in standard was used as a positive control. The raw sequence data for this effort are deposited in the sequence read archive (SRA) of the National Center for Biotechnological Information (NCBI) under BioProject (PRJNA956891).

### 2.3. Sequence analysis and statistical analysis

Reads from amplicon sequencing were trimmed to remove the PCR primers using Cutadapt v5.0 [16] and reads of less than 210 bases were removed from further analysis using Trimmomatic v0.39 [17]. Filtered reads were processed using Mothur v148.2 [18, 19]. to assemble, screen, and align contiguous sequences, call OTUs/ amplicon sequence variants (ASVs), make taxonomic assignments using the Silva nr database version 138.1 [20], and calculate the inverse Simpson Index. Samples with a proportion of *Corynebacterium* reads within one standard deviation of the mean for the extraction controls and field blanks were discarded from further analysis. The inverse Simpson index was based upon classification of contigs into ASVs using a similarity cutoff of 0.03. NMDS was performed using proportional abundance collapsed to the genus level of classification. The R package Vegan [21]was used to calculate the Bray-Curtis pairwise distance metric and perform non-metric multidimensional scaling (NMDS) analysis, PERMANOVA, beta dispersion, canonical analysis of principal components (CAPscale), and variance partitioning.

### 2.4. Quantification of PFOS sorption to isolates

Liquid cultures of R2A medium were inoculated from R2A plates and incubated overnight at 30°C. The grown cultures were then diluted 1:50 in 5 ml fresh R2A medium with 5 ppm PFOS. Cultures were then incubated at 30°C for 2 days. After incubation, 0.6 mL of the cultures were centrifuged, and 0.5 mL of the supernatant was diluted in 4 mL of 100% methanol. For whole culture extractions, the centrifugation step was omitted and 4 ml of 100% methanol was added to 0.5 ml of whole cell culture. Both sets of samples were stored at -20°C until sample cleanup.

High background in the medium interfered with direct measurements of PFAS, and required development of a new protocol to remove interfering compounds. Samples were cleaned and concentrated using solid phase extraction columns (Phenomenex, Torrance, CA, USA). Columns were conditioned using 10 ml methanol followed by 10 ml water. The whole sample was then added, and slowly passed through the column at a rate of less than 5 ml/min. After the sample was loaded, the sample containers and column were rinsed with 10 ml water followed by 1 ml of methanol, and dried by pulling air through for 5 minutes. Next, the columns were eluted with 10 ml of 2% ammonium hydroxide in methanol. The methanol was fully evaporated in a vacuum centrifuge and resuspended in 1 ml of 100% methanol.

Resuspended samples were analyzed by liquid chromatography – mass spectrometry (LC-MS). A Thermo-Scientific Vanquish LC system was set up for the analysis of PFAS containing samples. Samples were diluted 10:1 in LC-grade water, with a 25 µL injection volume. The primary separation column (Gemini 3 um C18, 100 x 3 cm) was installed along with a delay column (Luna Omega 5 um PS C18, 50 x 4.6 mm) installed upstream of the sample injection in line with the solvent introduction for the purpose of reducing possible PFAS contamination from the solvent pump system. The mobile phase gradient consisted of 5 mM ammonium acetate in water (solvent A) and 2 mM ammonium acetate in methanol (solvent B) as shown in table 1. The solvent flow rate was 0.7 ml per minute. A SCIEX 5600 MS was used in TOF scan mode to allow for the analysis of PFAS mixtures as well as detection of possible derivative products. Software controlled parameters were as follows: Curtain Gas = 30; Source Gas 1 = 40; Source Gas 2 = 60; IS Voltage = -4500; Temp = 450; Declustering Potential = -30; Collision Energy = -20.

**Table 1:** LC parameters for PFAS analysis.

| Gradient: | Time (min) | %B |
| --- | --- | --- |
|  | 0 | 55 |
|  | 1 | 55 |
|  | 5 | 65 |
|  | 8 | 95 |
|  | 8.5 | 99 |
|  | 12.95 | 99 |
|  | 13 | 10 |
|  | 15 | 10 |

### 2.5 Measurement of PFOS impact on bacterial growth

Strains were inoculated from an R2A plate into R2A liquid medium and grown overnight at 30C. The next morning, 10 µL of grown culture was added to 0.5 mL fresh R2A medium in 48 well clear bottomed plates, with or without 50 ppm PFOS, with 4 replicates of each condition. Plates were incubated at 30°C with shaking in a plate reader (Tecan, Männedorf, Switzerland) for 18 hours, with the optical density at 600 nm read every 10 minutes. The growth rate µ was calculated from the exponential phase of the growth curve using the R package *growthrates* v. 0.8.4 [22] with the all_easylinear tool, with h=8 and quota = 0.95, ignoring outlier datapoints that were >10% off their expected value based upon surrounding values. Growth rates were compared between the PFOS exposed cultures and unexposed controls using a two-sided t-tests, with a homoscedastic distribution.

## 3. Results

### 3.1. Sediment characterization

The sediment below 1.5 m was mostly sandy with some silty layers above 3.3 m, and some small patches of clay. By 7.2 m below the surface, it had more silt and clay, becoming sticky at times, and by 8.7 m, the sediment had become a dense clay. A layer rich in shell fragments was found just above the clay layer at the bottom of the cores, but shells were not seen anywhere else in the cores. The background color of the cores was a light gray-brown sand with light brown silty layers from 1.5 - 3 m, becoming medium gray by 3.3 m. Below 7.8 m, the gray sand was mixed with dark gray clay. The cores in the barrier were stained dark black by the CAC starting at 1.5 m, with some brown areas where the silty layers reduced penetration of the CAC between 1.5 - 3 m. The sand became lighter gray below 6.6 m, with a small amount of CAC that bled to this level and a few streaks of CAC evident down to 7.5 m.

### 3.2. Microbial community composition of PFAS impacted site

The microbial community composition of these cores was investigated using 16S amplicon sequencing. These samples did not yield much DNA, indicating that the microbial community did not grow to a high cell abundance. The low biomass led to us discarding 11 samples from the final analysis, because it was likely that many or even most of the non-*Corynebacterium* reads they contained were from the same source as the extraction controls and field blanks. These discarded samples were mostly in the 4.9, 5.8, and 7.3 m samples, indicating that there may have been a minimum of biomass in these middle depths. Interestingly, none of the 8.8 m samples were discarded, so it is possible that biomass increased below 7.3 m. As seen with other microbial communities, a large number of taxa were identified in these samples. Prior to further analysis, the OTU dataset was curated to remove OTUs that were found in less than seven of the samples to remove data that was likely to be sequencing noise. This left a total of 2054 OTUs. The total number of OTUs present in each sample ranged from 153 to 1070.

The inverse Simpson index was highly variable when compared by either depth or sample location (upstream – US; in barrier – IN; or downstream – DS) (Figure 1). Diversity was highest at 2.7 m and 3.4 m, but was significantly lower at 7.3 and 8.8 m, with t-test p-values of < 10-4 comparing between these depths (Fig. 1a). The DS samples did have a slightly higher diversity (Fig. 1b), but it was not statistically significant and may have simply been due to natural variability and the lower number of DS samples.

### 3.3. Community characteristics compared by location relative to barrier

NMDS was used to reduce the dimensionality of the data and enable visualization of the relationships between the microbial communities (Fig. 2a). While there was a high degree of variability between individual samples, the communities had a clear trajectory by depth in the NMDS plot. Some of the samples grouped together by depth, especially depths 2.7 m, 7.3 m, and 8.8 m, but the intermediate depths, 3.4 m to 5.8 m were more variable. There was a small amount of separation by the year the samples were obtained. The samples from 2022, shortly after the CAC was injected to form the barrier, tend to be closer to the bottom of the NMDS plot. This effect is less pronounced at 7.3 and 8.8 m. These samples were below the level of the barrier, with only small amounts of visible CAC.

**Fig. 2.**
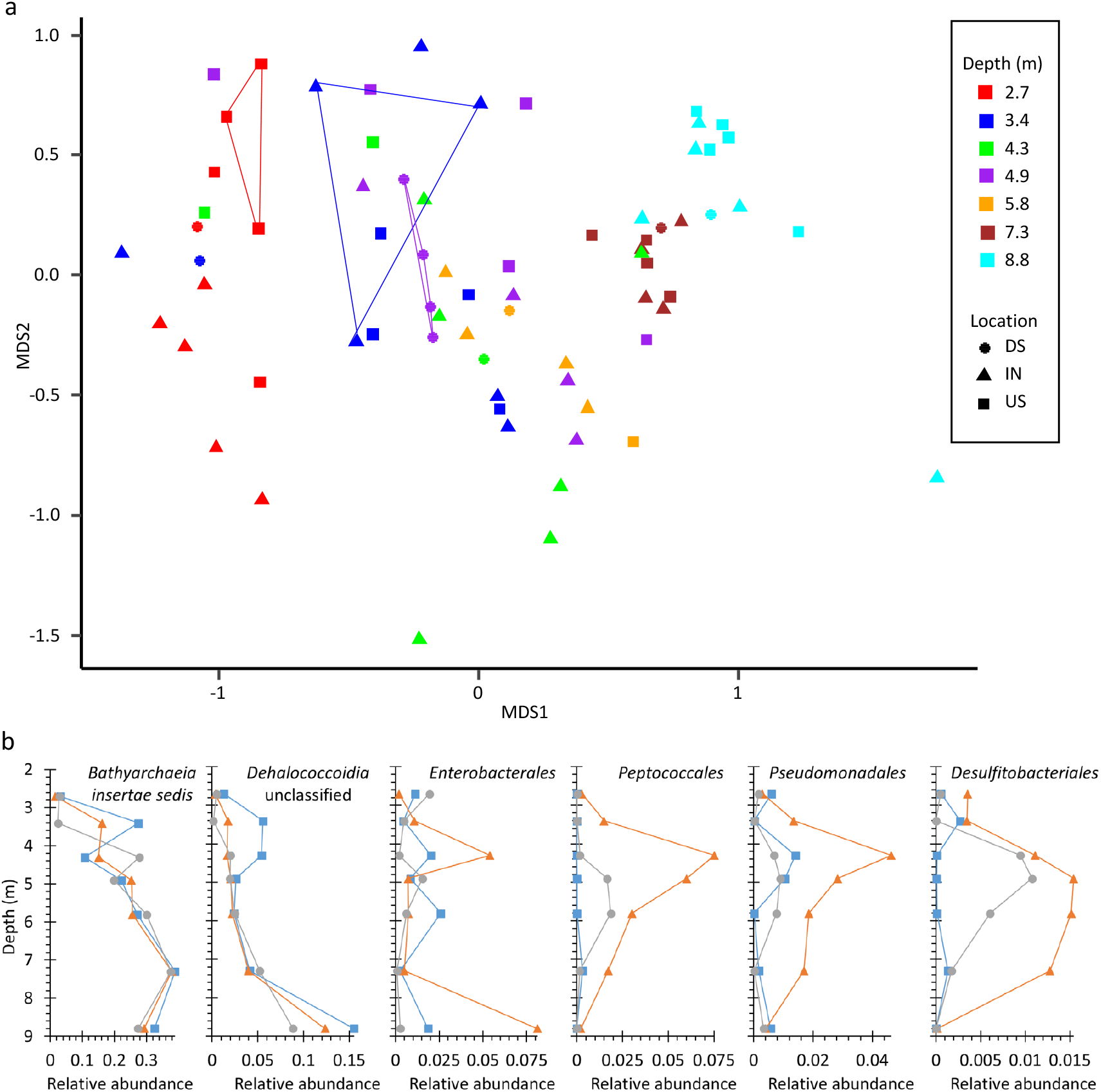
Ordination plots of samples at the genus level. (a) Nonmetric multidimensional scaling of Bray-Curtis distance between samples was used to depict relationships between the microbial communities. Color of the points indicates the depth for that sample, while the shape indicates the location or the core relative to the CAC barrier (square = upstream, triangle = in barrier, circle = downstream of barrier). Convex hulls are depicted around replicate samples. (b) Relative abundance of select orders at depth by location. Note the x-axes are scaled by the maximum relative abundance for each order

Variance partitioning quantified the role that the sample attributes played in influencing differences between the samples. It indicated that depth was responsible for the majority (21.26%) of variability that could be attributed to a metadata characteristic, with the year of sampling contributing 2.6%, of variability, and only 0.75% coming from the location relative to the barrier. This may be an underestimate, however, because this did not account for the fact that 7.3 m and 8.8 m should have minimal impact from the CAC barrier. There was still a high residual (75.4%), likely a result of the high diversity of the microbial community present in these samples and the patchy and highly variable nature of sediment microbial communities.

Replicate 10-15 g samples were collected from three core sections to provide context for how centimeter-scale differences in sampling location affected variability. While there were clear differences in the abundance of OTUs within the three replicate samples, the Bray-Curtis dissimilarity metric was smaller between replicates than between other the samples from the same depth. These differences were significant for the IN6 3.4 m and DS9 4.9 m replicates (t-test p-values = 0.016 and 4.7 x 10-9, respectively) but not for the US5 2.7 samples (p-value = 0.17). Variability between replicates did not appear to be related to extraction batch, because two of the replicate samples that were most different by NMDS (IN6 3.4A and IN6 3.4B) were extracted in the same batch, but this was not rigorously tested.

### 3.4. Community composition and role of microbial taxa identified

A large number of the microbes that were identified here are unclassified, which creates difficulty in assigning roles based upon taxonomic identity. One of the most abundant bacterial phyla was the *Bacillota*, which includes *B. subtilis*, but the order *Bacillales* was not very abundant. However, the *Bacillota* orders *Peptococcales, Desulfitobacterales*, and unclassified *Clostridia* (all typically obligate anaerobes) were abundant. The *Peptococcales* and *Desulfitobacterales*, made up a larger proportion of the community in the CAC treated samples than in the upstream samples, and had slightly higher relative abundance in the downstream samples (Fig. 2b). The genera that were driving this increased relative abundance were *Desulfitispora, Desulfosporosinus*, and an unclassified genus within the *Desulfitobacteriaceae*. The phylum *Bathyarchaeota* was one of the most abundant phyla, including the order “*Bathyarchaeia* insertae sedis". These Archaea are common inhabitants of subsurface sediments and are predicted to have a diverse set of metabolic properties that makes them difficult to assign a role [23, 24]. We see a trend in increased relative abundance of “*Bathyarchaeia* insertae sedis” with depth and a decrease in “Archaea unclassified". Members of the phylum *Chloroflexota*, including the class *Dehalococcoidia* known for dehalogenation, were more abundant at the lower levels of the cores, but no isolates of this phylum were obtained.

Many of the isolates reported below belonged to orders found within the phylum *Pseudomonadota* (formerly known as *Proteobacteria*). Despite their ease of culture, they made up a relatively small fraction of the environmental community, with only 7.3% of the relative abundance. This was mostly made up of four orders, the *Enterobacterales* (isolates *Serratia* sp. and *Klebsiella* sp.), *Pseudomonadales (*isolates *Pseudomonas* sp. CRBF and *Acinetobacter* sp.), *Burkholderiales* (*Massilia* sp.). The *Micrococcales* include the genus *Paenarthrobacter* from which an isolate was recovered, but the majority of sequences identified from this order belonged to unclassified genera within the families *Micrococcaceae* and *Microbacteriaceae* as well as unclassified families within the order *Micrococcales*.

The order *Flavobacteriales*, which contains the genus *Chryseobacterium*, was identified in most samples, but only had a relative abundance greater than 1% in one of the samples.

### 3.5. PFOS uptake by bacterial cultures

Ten bacterial strains (Table 2) were used as representative bacteria to quantify the amount of PFOS that is sorbed to bacterial cells (Fig. 3a). Six were isolated from these PFAS impacted sediment cores, one strain from an enrichment culture, and three strains were obtained from a culture collection. The PFOS sorption by each strain was calculated as the difference between the PFOS concentration in the abiotic controls and the culture supernatant divided by the amount recovered in the abiotic control (Fig. 3a). PFOS can adsorb to many labware surfaces, and is also present as a contaminant in some labware, which necessitated abiotic controls to account for potential differences in PFOS recovery during cleanup, as well as a PFOS free control to check for contamination. These were used to set a baseline for how much PFOS could be recovered in the absence of bacteria. To verify that PFOS was sorbing to cellular material and not being lost to labware or extraction, recovery from whole cells in several strains was examined (Fig. 3b). Recovery of PFOS from methanol extracted samples of the whole culture medium including cells was similar to the abiotic control (close to 100%), while extractions of the supernatant of those cultures were lower and similar to supernatant values for previous cultures. This indicated that the PFOS missing from the supernatant in Fig. 3a is bound up by the bacterial cells, and that the PFOS was not altered by the cells. No fluoride was detected, and no other products were detected via LC-MS. The pH of the bacterial cultures after growth did not have any correlation to PFOS sorption (Table 2).

**Table 2.** Bacterial strains used. pH was measured after 2 days growth in R2A.

| Organism | pH | Source |
| --- | --- | --- |
| *Klebsiella* sp. T6R-B | 7.7 | This study |
| *Chryseobacterium* sp. CBR-D | 7.5 | This study |
| *Serratia* sp. U2P | 7.7 | This study |
| *Paenarthrobacter* sp. CBR-A | 7.7 | This study |
| *Acinetobacter* sp. B1 | 8.6 | This study |
| *Pseudomonas* sp. CBR-F | 7.8 | [13] |
| *Serratia* sp. T2R-B1 | 7.8 | [25] |
| *Escherichia coli* K12 (C3000) | 7.0 | ATCC 15597 |
| *Pseudomonas fluorescens* | 7.5 | ATCC 13525 |
| *Bacillus subtilis* | 7.9 | ATCC 23857 |

**Fig. 3.**
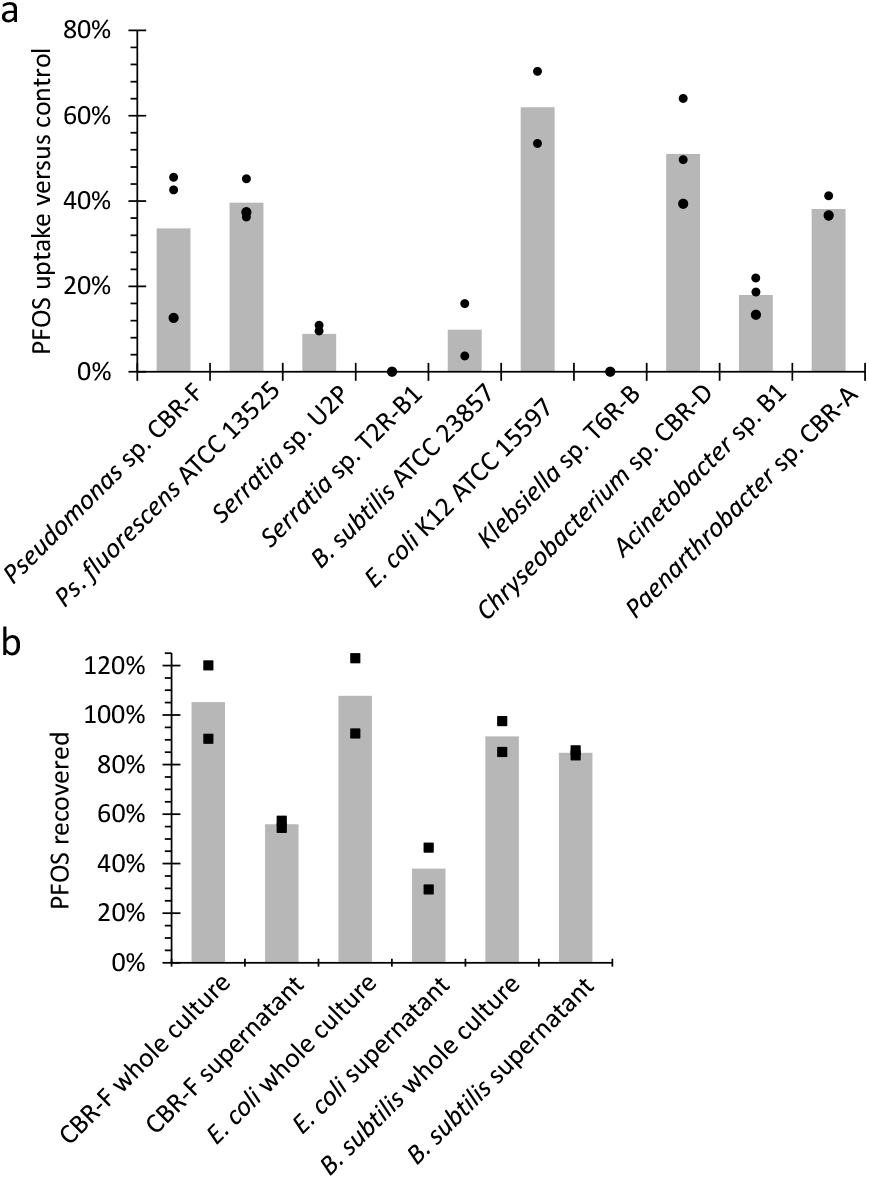
PFOS sorption by bacterial cells. a) Percentage of PFOS removed from medium by bacterial cultures after 2 days of growth. Percentages were calculated by subtracting the PFOS detected in the supernatant from the PFOS detected in the abiotic control. b) Extraction of select whole culture samples with methanol, normalized to abiotic control

### 3.6. Physiological characteristics that may influence PFOS sorption

Next, we determined whether the sorption of PFOS influenced the growth of the bacteria. We grew each strain with and without 50 ppm PFOS (Fig. 4). PFOS reduced the growth rate of six of these strains, but did not change in the timing or length of the growth phases of most of the cultures (Fig. 4a-c). The growth rate of *Bacillus subtilis* was significantly lower in the presence of PFOS, during the exponential phase of growth, but this strain entered a phase with linear growth where its optical density continued to increase at later time points, which may indicate changes in cell size or shape. Three other strains did exhibit diauxic growth, with a substantial lag phase for *E. coli, P. fluorescens, and Paenarthrobacter* sp. CBR-A between an initial logarithmic phase and a secondary logarithmic phase.

**Fig. 4.**
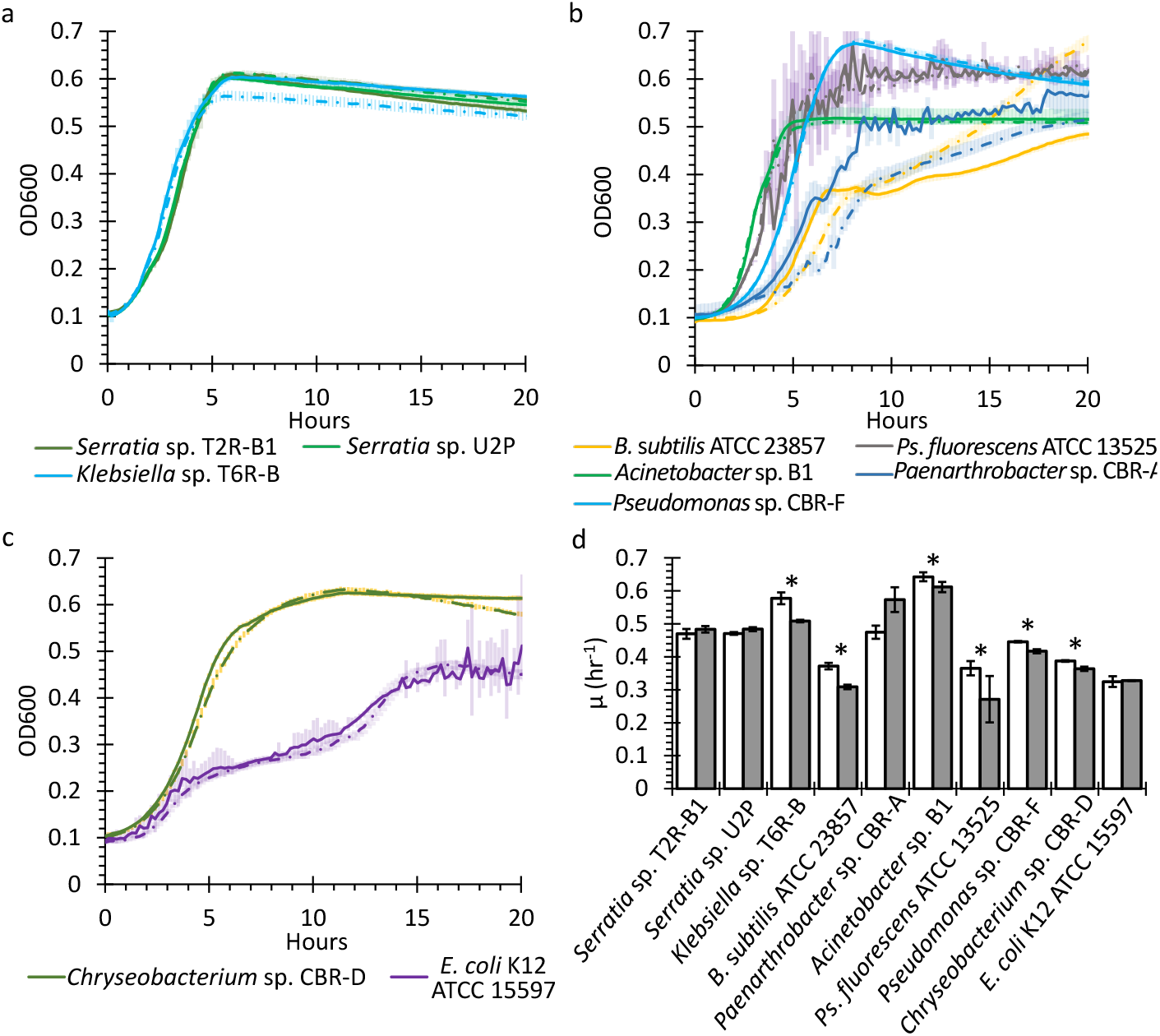
Growth of strains in R2A medium with and without PFOS. Growth curves are grouped by PFOS sorption (a) low sorption, (b) medium sorption, and (c) high sorption. Solid lines show the mean optical density at 600 nm (OD600) without PFOS while dotted lines show OD600 with 50 ppm PFOS. (d) Growth rates calculated from the maximum logarithmic portion of the growth curve for cultures without (open bars) and with PFOS (filled bars). *p-value with two-sided t-test < 0.05. The *Paenarthobacter* sp. CBR-A growth curve data was unreliable, and the high growth rate for the PFOS exposed culture is likely an artifact of excessive noise

This is likely a result of the low concentrations of each component of R2A medium, leading to a rapid depletion. However, there was no apparent relationship between PFOS sorption and change in growth rate. The two strains with the highest PFOS uptake, *E. coli* and *Chryseobacterium* sp. CBR-D, did not show any change in growth between the PFOS and control.

## 4. Discussion

### 4.1. Subsurface microbial community

The subsurface microbial community was very diverse, with 2054 OTUs found in at least seven of the samples. As expected, the diversity was higher at the shallower depths, and dropped to a low at 7.2 m below the surface. This may be reflective of the hydrology and geology of the site, because a thick layer of clay starting around 8.7 m below the surface forms a confining layer and keeps the water table high in this area, leading to stable conditions. Closer to the surface, it is likely that the community is exposed to a more variable regime, for example, during the 2022 sampling the core sections from 3-4.5 m were noted as being dry to damp, while in 2024 it was quite wet. While the diversity was not significantly different between the US, IN, and DS samples, there were still differences in the relative abundance of some taxonomic groups between these samples.

The community had a large fraction of poorly or uncharacterized bacteria and archaea, which highlights understudied nature of this environment. The relative abundance of Archaea increased with depth, as has been previously reported [26], with much of this shift driven by the *Bathyarchaeia insertae sedis*. Within the Archaea, *Bathyarchaeia insertae sedis* was the order level taxon with the highest relative abundance. The *Bathyarchaeia* are known for their metabolic flexibility and widespread distribution in sediments, making it difficult to assign a function to their presence here. It is difficult to assign a putative function to most of the Archaeal members of this community, but one relatively common taxon was the class *Nitrososphaeria*, which includes the order level taxa Group 1.1c, *Nitrososphaeria*_unclassified, *Nitrososphaerales*, and *Nitrosotaleales*. This class is known for autotrophic ammonia oxidizers, although it has also been noted that they are capable of methane oxidation as well [27, 28].

Among the bacteria, the relative abundance of the *Peptococcales* (primarily *Desulfitispora* spp.) and *Desulfitobacterales* peaked in the middle depths (3.4 – 5.8 m below surface) which is the depth range with the highest amount of CAC. The samples from 2022, shortly after CAC injection had the higher abundance, and these taxa were rare upstream of injection sites and less prevalent downstream. This suggest that they are associated with the presence of the CAC barrier, although it is unclear why.

Three possible explanations are that an additive in the injection mix served as a substrate that enriched for these two groups of bacteria in particular, there was a high abundance of *Peptococcaceae* spores or vegetative cells in the injection mixture, or something about the CAC itself served to enrich for these bacteria. A number of recent attempts to reorganize the *Peptococcales* resulted in splitting off the *Desulfitobacterales* that were previously included within this family. Several *Desulfitobacterium* species have been described as dehalogenating organohalides, including D. *hafniens*e, *D. dehalogenans*, and *D. chlororespirans* [29-31] and a number of other isolates, enrichments, and metagenome amplified genomes have been identified as having some affinity to the *Peptococcales* and its former members [32, 33]. It is likely that the shallower samples had a low relative abundance of the anaerobic *Peptococcaceae* due to higher oxygen concentrations in near the surface of the water table.

With such low biomass, several of the PCRs did not yield enough good data to include in this analysis, but the samples that were included here do show that they are of sufficient quality to enable interpretation. Samples that were expected to be more similar to each other (the deep samples from all cores and the replicates from the same core and depth) were indeed mostly similar, although a few of the samples appeared to group with samples of other depths and locations in the NMDS plot. Some of these included US5 3.4 m and DS9 3.4 m, and US5 4.9 m, but these may simply be a result of contamination from water drainage within the bore hole.

The three sets of replicate samples set an upper bound on the variability due to sampling. The depths for replicate samples (2.7 m, 3.4 m, 4.9 m) were selected without knowing that these depths had a higher inherent variability, but because these samples had the highest variability, the replicates represent an upper bound of variation. The 2.7 m and 4.9 m replicates were tightly grouped along the NMDS1 axis, but had more variability on the NMDS2 axis. A convex hull drawn around these points did not include any other samples. However, the IN6 3.4 m replicates were much more variable along both axes, and a hull drawn around these three points included five others. The 3.4 m samples were the most variable of any depth, so it is possible that this is more indicative of this being a transitional depth between the shallow more surface influenced sediments and the deeper sediments.

### 4.2. Sorption of PFOS and physiology of bacterial isolates

Sorption of PFOS by *Pseudomonas* species and *Bacillus subtilis* in the present work were comparable to results reported previously [9-11], with both strains of *Pseudomonas* and *Bacillus subtilis* sorbing PFOS. In contrast to the results of Butzen et al [11], we observed sorption at a pH close to neutral for *Pseudomonas*. This may have resulted from differences in protocols – we used rich medium over two days in our experiments while Butzen et al used washed cells for a brief period in their method. It is also possible that the species of *Pseudomonas* (*P. putida* in their work vs *P. fluorescens* and new isolates in ours) affected the results. Our new strains, displayed an unexpected result: two strains belonging to the genus *Serratia* and one belonging to the genus *Klebsiella* showed little to no sorption of PFOS. Both these genera belong to the *Enterobacteriaceae* family – however, *E. coli* K12, belonging to the same family, displayed the highest sorption of any bacterium. All of the strains grew in the presence of 50 ppm PFOS, although about half of them exhibited slower growth (Fig. 4d), but this was not correlated to the level of PFOS sorbed by the strain. It is interesting to note the effect of PFOS on the growth of *Pseudomonas sp*. CBR-F was less than that of *Pseudomonas fluorescens* in the presence of PFOS. This difference may indicate that *Pseudomonas sp*. CBR-F may have evolved some resistance to PFAS due to its residence in a PFAS impacted environment.

### 4.3. Potential Impact of differential sorption of PFOS

We compared the isolates to the microbial community of the cores from which they were isolated. While the most abundant orders in the environment were not isolated from these cores, we were able to obtain isolates from orders are relatively abundant in our sequencing data set. The isolates mostly did not belong to the most prevalent genera within those orders but can be expected to be physiologically similar. Despite not being able to identify a clear trend of sorption with respect to fatty acid composition, there other physiological features that may be correlated with the PFAS sorption.

These include surface charge, exopolysaccharide composition, and specific surface proteins. While investigating the mechanism behind differential sorption was beyond the scope of the present work, it might represent an interesting future avenue of research, especially since these can all vary in response to specific environmental conditions [34-36]. Recent work by Lindell et al [37] found that perfluorononanoic acid is accumulated intracellularly in *E. coli*, but that it is actively transported out of the cell, leading to the possibility that adsorption, absorption, and active transport all play a role in PFAS accumulation by microbes.

The microbial community that we investigated here represents sediment associated biomass within the aquifer as opposed to many previous studies that used groundwater as the starting material [8, 38]. This gives a unique perspective on the potential role of these microbes in binding PFOS, because these microbes may be more sedentary. High PFOS binding microbes within this community may contribute to slowing down the transport of PFOS within the groundwater. On the other hand, free living microorganisms may contribute to the transport of PFOS and other PFAS in the subsurface environment by sorbing it and preventing its immobilization. The specific makeup of a local microbial community may therefore have a significant impact on the transport rate of PFOS in the environment, but at present we do not have enough information to accurately assess whether a given community will have a net-positive or net-negative effect on this rate.

## 5 Conclusions

Understanding the migration of PFOS through soil, sediment, and groundwater is important to management of these contaminants in groundwater and agricultural soils. The mobility of PFOS through these matrices is complex, with particle size and type, presence and type of plants, and rate of water movement all contributing to its fate. The wide range of PFOS sorption means that microbial interactions are another factor that needs consideration when developing models for the environmental mobility of PFAS. The diverse microbial community in the subsurface may include microbes that can bind PFAS and form long-lived biofilms, thus retaining the PFAS. In the case of free-living cells with a high affinity for PFAS, they may enhance PFAS mobility by preventing it from binding soil and sediment particles. The present work only investigated PFOS, and previous work has only investigated the affinity of microbes for five additional PFAS chemicals [9, 11], leading to uncertainty in microbial-PFAS interactions. The present work, and previous investigations [9, 11] of PFAS sorption have focused on aerobic bacteria that are not particularly representative of the native organisms. Further investigations with anaerobic microbes, especially Archaea, are needed, considering the variation between isolates from the same genus. The subsurface microbial community consisted of taxa that are often associated with sediment environments, but there were indication that presence of PFAS or the CAC barrier used to treat it affected the microbial community. The abundance of *Peptococcales* may be a result of the CAC injections, as their relative abundance was significantly higher shortly after injections and was higher in the samples with the greatest exposure to the CAC, but no causative relationship could be established, so further research is needed to understand whether their growth was stimulated by the CAC, or as a side effect of the injection process. A better understanding of the microbial attributes that lead to PFOS sorption to bacterial cells and a better understanding of the microbial communities that form under different conditions will allow modeling of microbial interactions with PFOS and other PFAS when predicting the environmental transport of PFAS in groundwater systems.

## Acknowledgements

The authors would like to thank Graig Lavorgna, Paul Hatzinger, and Sara Foxwell from APTIM, and Keith Gaskill from Regenesis for allowing us to take samples of their cores, and Anthony Danko from

U.S. Naval Facilities Engineering and Expeditionary Warfare Center (NAVFAC EXWC) for facilitating site access. This work was supported through the U.S. Office of Naval Research via U.S. Naval Research Laboratory base program WU 991AE3.

## References

1 Anderson RH, Thompson T, Stroo HF, Leeson A (2021) US Department of Defense–Funded Fate and Transport Research on Per-and Polyfluoroalkyl Substances at Aqueous Film–Forming Foam–Impacted Sites: US Department of Defense PFAS fate and transport research. Environ Toxicol Chem 40(1):37–43

2 Brusseau ML, Anderson RH, Guo B (2020) PFAS concentrations in soils: Background levels versus contaminated sites. Sci Total Environ 740:140017

3 Environmental Protection Agency Title 40 Chapter I Subchapter D Part 141 National Primary Drinking Water Regulaons. htps://www.ecfr.gov/current/tle-40/part-141. Accessed 8/18/2025, 2025

4 LaFond JA, Hatzinger PB, Guelfo JL, Millerick K, Jackson WA (2023) Bacterial transformation of per- and poly-fluoroalkyl substances: a review for the field of bioremediation. Environmental Science: Advances 2(8):1019–1041

5 Xu R, Tao W, Lin H, Huang D, Su P, Gao P, Sun X, Yang Z, Sun W (2022) Effects of perfluorooctanoic acid (PFOA) and perfluorooctane sulfonic acid (PFOS) on soil microbial community. Microb Ecol 83(4):929–941

6 Ke Y, Chen J, Hu X, Tong T, Huang J, Xie S (2020) Emerging perfluoroalkyl substance impacts soil microbial community and ammonia oxidation. Environ Pollut 257:113615

7 Weathers TS, Harding-Marjanovic K, Higgins CP, Alvarez-Cohen L, Sharp JO (2016) Perfluoroalkyl acids inhibit reductive dechlorination of trichloroethene by repressing Dehalococcoides. Environ Sci Technol 50(1):240–248

8 O’Carroll DM, Jeffries TC, Lee MJ, L. ST, Yeung A, Wallace S, Batye N, Patch DJ, Manefield MJ, Weber KP (2020) Developing a roadmap to determine per-and polyfluoroalkyl substances-microbial population interactions. Sci Total Environ 712:135994

9 Fitzgerald NJ, Wargenau A, Sorenson C, Pedersen J, Tufenkji N, Novak PJ, Simcik MF (2018) Partitioning and accumulation of perfluoroalkyl substances in model lipid bilayers and bacteria. Environ Sci Technol 52(18):10433–10440

10 Naumann A, Alesio J, Poonia M, Bothun GD (2022) PFAS fluidize synthetic and bacterial lipid monolayers based on hydrophobicity and lipid charge. Journal of environmental chemical engineering 10(2):107351

11 Butzen ML, Wilkinson JT, McGuinness SR, Amezquita S, Peaslee GF, Fein JB (2020) Sorption and desorption behavior of PFOS and PFOA onto a Gram-positive and a Gram-negative bacterial species measured using parcle-induced gamma-ray emission (PIGE) spectroscopy. Chemical Geology 552:119778

12 Ji B, Zhao Y (2024) Interactions between biofilms and PFASs in aquatic ecosystems: literature exploration. Sci Total Environ 906:167469

13 Baker IR, Colston SM, Hervey J, Eddie BJ, Bird LJ (2025) Complete genome of a fluorescent Pseudomonas sp. isolated from a PFAS groundwater treatment site. Microbiology Resource Announcements 14(2):e00976–00924

14 Apprill A, McNally S, Parsons R, Weber L (2015) Minor revision to V4 region SSU rRNA 806R gene primer greatly increases detection of SAR11 bacterioplankton. Aquat Microb Ecol 75(2):129–137

15 Parada AE, Needham DM, Fuhrman JA (2016) Every base maters: assessing small subunit rRNA primers for marine microbiomes with mock communities, me series and global field samples. Environ Microbiol 18(5):1403–1414

16 Martin M (2011) Cutadapt removes adapter sequences from high-throughput sequencing reads. EMBnet journal 17(1):10–12

17 Bolger AM, Lohse M, Usadel B (2014) Trimmomatic: a flexible trimmer for Illumina sequence data. Bioinformatics 30(15):2114–2120

18 Kozich JJ, Westcot SL, Baxter NT, Highlander SK, Schloss PD (2013) Development of a dual-index sequencing strategy and curation pipeline for analyzing amplicon sequence data on the MiSeq Illumina sequencing platform. Appl Environ Microbiol 79(17):5112–5120

19 Schloss PD, Westcot SL, Ryabin T, Hall JR, Hartmann M, Hollister EB, Lesniewski RA, Oakley BB, Parks DH, Robinson CJ (2009) Introducing mothur: open-source, platform-independent, community-supported software for describing and comparing microbial communties. Appl Environ Microbiol 75(23):7537–7541

20 Quast C, Pruesse E, Yilmaz P, Gerken J, Schweer T, Yarza P, Peplies J, Glöckner FO (2012) The SILVA ribosomal RNA gene database project: improved data processing and web-based tools. Nucleic Acids Res 41(D1):D590-D596

21 Oksanen J, Simpson G, Blanchet F, al. e (2024) Vegan: Community Ecology Package (R Package Version 2.6-8). 2024. In: R package version 2.6-8, htps://CRAN.R-project.org/package=vegan ed.,

22 Petzoldt T (2019) Estimation of growth rates with package growthrates. In: R package version 0.8.4, htps://CRAN.R-project.org/package=growthrates ed.,

23 Meng J, Xu J, Qin D, He Y, Xiao X, Wang F (2014) Genetic and functional properties of uncultivated MCG archaea assessed by metagenome and gene expression analyses. The ISME journal 8(3):650–659

24 Lloyd KG, Schreiber L, Petersen DG, Kjeldsen KU, Lever MA, Steen AD, Stepanauskas R, Richter M, Kleindienst S, Lenk S (2013) Predominant archaea in marine sediments degrade detrital proteins. Nature 496(7444):215–218

25 Baker IR, Colston SM, Hervey WJ, Bird LJ (2023) Complete genome of a Serratia species isolated from PFAS-impacted soil. Microbiology Resource Announcements 12(12):e00640–00623

26 Turner S, Mikuta R, Meyer-Stüve S, Guggenberger G, Schaarschmidt F, Lazar CS, Dohrmann R, Schippers A (2017) Microbial community dynamics in soil depth profiles over 120,000 years of ecosystem development. Frontiers in microbiology 8:874

27 Pester M, Schleper C, Wagner M (2011) The Thaumarchaeota: an emerging view of their phylogeny and ecophysiology. Curr Opin Microbiol 14(3):300–306

28 Könneke M, Schubert DM, Brown PC, Hügler M, Standfest S, Schwander T, Schada von Borzyskowski L, Erb TJ, Stahl DA, Berg IA (2014) Ammonia-oxidizing archaea use the most energy-efficient aerobic pathway for CO2 fixation. Proceedings of the National Academy of Sciences 111(22):8239–8244

29 Sanford RA, Cole JR, Löffler FE, Tiedje JM (1996) Characterization of Desulfitobacterium chlororespirans sp. nov., which grows by coupling the oxidation of lactate to the reductive dechlorination of 3-chloro-4-hydroxybenzoate. Appl Environ Microbiol 62(10):3800–3808

30 Chrisansen N, Ahring BK (1996) Desulfitobacterium hafniense sp. nov., an anaerobic, reducvtiely dechlorinang bacterium. Int J Syst Evol Microbiol 46(2):442–448

31 Utkin I, Woese C, Wiegel J (1994) Isolation and characterization of Desulfitobacterium dehalogenans gen. nov., sp. nov., an anaerobic bacterium which reducvtiely dechlorinates chlorophenolic compounds. Int J Syst Evol Microbiol 44(4):612–619

32 Holland SI, Ertan H, Montgomery K, Manefield MJ, Lee M (2021) Novel dichloromethane-fermenting bacteria in the Peptococcaceae family. The ISME journal 15(6):1709–1721

33 Kleindienst S, Higgins SA, Tsementzi D, Chen G, Konstantinidis KT, Mack EE, Löffer FE (2017) ‘Candidatus Dichloromethanomonas elyunquensis’ gen. nov., sp. nov., a dichloromethane-degrading anaerobe of the Peptococcaceae family. Syst Appl Microbiol 40(3):150–159

34 Blanco Y, Rivas LA, González-Toril E, Ruiz-Bermejo M, Moreno-Paz M, Parro V, Palacín A, Aguilera Á, Puente-Sánchez F (2019) Environmental parameters, and not phylogeny, determine the composition of extracellular polymeric substances in microbial mats from extreme environments. Sci Total Environ 650:384–393

35 Poortinga AT, Bos R, Norde W, Busscher HJ (2002) Electric double layer interactions in bacterial adhesion to surfaces. Surface science reports 47(1):1–32

36 Collins YE, Stotzky G (1992) Heavy metals alter the electrokinetic properties of bacteria, yeasts, and clay minerals. Appl Environ Microbiol 58(5):1592–1600

37 Lindell AE, Grießhammer A, Michaelis L, Papagiannidis D, Ochner H, Kamrad S, Guan R, Blasche S, Ventimiglia LN, Ramachandran B (2025) Human gut bacteria bioaccumulate per-and polyfluoroalkyl substances. Nature Microbiology:1–18

38 Tang Z, Song X, Xu M, Yao J, Ali M, Wang Q, Zeng J, Ding X, Wang C, Zhang Z (2022) Effects of co-occurrence of PFASs and chlorinated aliphatic hydrocarbons on microbial communities in groundwater: a field study. J Hazard Mater 435:128969

